# The Macromolecular MR Spectrum in Healthy Aging

**DOI:** 10.1101/2021.08.27.457961

**Authors:** Steve C.N. Hui, Tao Gong, Helge J. Zöllner, Yulu Song, Saipavitra Murali-Manohar, Georg Oeltzschner, Mark Mikkelsen, Sofie Tapper, Yufan Chen, Muhammad G. Saleh, Eric C. Porges, Weibo Chen, Guangbin Wang, Richard A. E. Edden

**Affiliations:** The Russell H. Morgan Department of Radiology and Radiological Science, Johns Hopkins University School of Medicine, Baltimore, MD, USA; F.M. Kirby Research Center for Functional Brain Imaging, Kennedy Krieger Institute, Baltimore, MD, USA; Department of Radiology, Shandong Provincial Hospital, Cheeloo college of Medicine, Shandong University, Jinan, Shandong, China; Department of Radiology, Shandong Provincial Hospital Affiliated to Shandong First Medical University, Jinan, Shandong, China; Department of Diagnostic Radiology and Nuclear Medicine, University of Maryland School of Medicine, Baltimore, MD, USA; Center for Cognitive Aging and Memory, University of Florida, Gainesville, FL, USA; McKnight Brain Research Foundation, University of Florida, FL, USA; Department of Clinical and Health Psychology, University of Florida, Gainesville, FL, USA; Department of Radiology, Weill Cornell Medicine, New York, NY, USA; Philips Healthcare, Shanghai, China

**Keywords:** magnetic resonance spectroscopy, macromolecule, PRESS, pre-inversion pulse, healthy aging

## Abstract

**Purpose:** Mobile macromolecules (MMs) from amino acids, cytosolic proteins and mobile lipids contribute a significant spectral background underlying the metabolite signals in the MR spectrum. A recent consensus recommends that MM contributions should be removed or included in modeling basis sets for determination of metabolite concentrations and/or metabolite ratios. The purpose of this study was to acquire the MM spectrum from healthy participants at a range of ages, and to investigate changes in the signals with age and sex groups.

**Methods:** Inversion time (TI) series were acquired to determine an optimal inversion time to null the metabolite signals. Experiments were carried out using a single adiabatic hyperbolic-secant inversion pulse. After the preliminary experiment, 102 volunteers (49M/53F) between 20 and 69 years were recruited for in vivo data acquisition in the centrum semiovale (CSO) and posterior cingulate cortex (PCC). The protocol consisted of a T_1_-weighted MPRAGE for structural images, followed by PRESS localization using a voxel size of 30 × 26 × 26 mm^3^ with pre-inversion (TR/TI 2000/600 ms) and CHESS water suppression. Metabolite-nulled spectra were modeled using a reduced basis set (NAA, Cr, Cho, Glu) and a flexible spline baseline (0.1 ppm knot spacing) followed by subtraction of the modeled metabolite signals to yield a ‘clean’ MM spectrum, using the Osprey software. Pearson’s correlation coefficient was calculated between integrals and age for the 14 MM signals between 0.9–4.2 ppm. One-way ANOVA was performed to determine differences between age groups. An independent t-test was carried out to determine differences between sexes. Relationships between brain tissues with age and sex groups were also measured.

**Results:** MM spectra were successfully acquired in 99 (CSO) and 96 (PCC) of 102 subjects. No significant correlations were seen between age and MM integrals. One-way ANOVA also suggested no age-group differences for any MM peak (all p > 0.004). No differences were observed between sex groups. The voxels were segmented as 80 ± 4% white matter, 18 ± 4% gray matter, and 2 ± 1% CSF for CSO and 28 ± 4% white matter, 61 ± 4% gray matter and 11 ± 1% CSF for PCC. WM and GM showed a significant (p < 0.05) negative linear association with age in the WM-predominant CSO (R = −0.29) and GM-predominant PCC regions (R = −0.57) respectively while CSF increased significantly with age in both regions.

**Conclusion:** Our findings indicate that the MM spectrum is stable across a large age range and between sexes, suggesting a pre-defined MM basis function can be used for linear combination modeling of metabolite data from different age and sex groups.

**Highlights:**

1. A large publicly available MM-aging dataset is presented.
2. Macromolecule signals do not change with age between 20 and 70.
3. There is no sex difference for macromolecule integrals.

## 1. Introduction

Short-TE ^1^H magnetic resonance (MR) spectra acquired in the brain contain relatively narrow signals from metabolites and broader signals from higher-molecular-weight macromolecules (MMs). The MM signals originate from amino acids in cytosolic proteins, peptides and mobile lipids (Behar and Ogino, 1993; Behar et al., 1994; Cudalbu et al., 2021). Characterization of these MM signals is important both in its own right, to understand changes in the cellular microenvironment, and for reliable quantitative measurements of the metabolites (Cudalbu et al., 2021). It is possible to acquire an MM spectrum by pre-inversion at an inversion time (TI) corresponding to the null point of the metabolite signals, exploiting the different longitudinal relaxation times (T_1_) of metabolites and MMs. The T_1_s of the MM signals are approximately four times shorter than metabolite T_1_s, meaning that MM signals are almost fully relaxed at this metabolite-nulling TI (Xin et al., 2013).

Metabolite T_1_ can be measured by inversion recovery and saturation recovery (Taylor et al., 2016), with the former offering more dynamic range and the latter being more easily implemented. The literature on human brain T_1_ at 3T is somewhat limited (Ethofer et al., 2003; Mlynarik et al., 2001; Traber et al., 2004) especially for signals other than the three methyl singlets. It is notable that these non-methyl signals have generally shorter T_1_s than the methyl singlets, and are therefore likely to be positive at singlet-nulling TIs. There is, to our knowledge, only one study quantifying the T_1_ of all observable MM peaks at 3T, and they are between 225 and 400 ms (Hoefemann et al., 2020).

The broad range of metabolite T_1_s results in incomplete metabolite nulling in MM spectra, with e.g., Schaller et al. describing a process for addressing residual metabolite peaks (Schaller et al., 2013). Cr_3.0_ has the median methyl singlet T_1_, and it is often seen that residual NAA signals are positive and residual choline (Cho) signals are negative in macromolecular spectra. T_1_ contrast between residual metabolite signals is reduced at shorter repetition time (TR) (Mlynarik et al., 2001). Given the difficulties of differentiating broader coupled signals like mI from the macromolecular signal, it is beneficial to acquire the macromolecular signal at relatively short TR and relatively short TI anticipating negative residual signals for NAA, Cr_3.0_ and Glu.

Studies of metabolite concentration changes with normal and abnormal aging may be confounded by changes in metabolite relaxation or macromolecular signals (Hupfeld et al., 2021; Lind et al., 2020; Porges et al., 2017). For example, GABA+, which includes a co-edited MM contribution, has been reported to change across the lifespan (Aufhaus et al., 2013; Gao et al., 2013; Ghisleni et al., 2015; Porges et al., 2021; Porges et al., 2017; Saleh et al., 2020; Simmonite et al., 2019), but a recent study of MM-suppressed GABA MRS failed to find a relationship in a developmental cohort indicating that age-associated MM changes may have contributed to the previous reports of a developmental change (Bell et al., 2021). Transverse relaxation times of metabolites have been reported to decrease with age (Marjanska et al., 2013). A recent consensus paper (Cudalbu et al., 2021) suggested that the macromolecule spectrum is different in older vs. younger adults. However, this was based upon two studies of small populations: the first study included only 4 young and 3 older subjects (Marjanska et al., 2018) and the second 7 younger (<25 years), 23 mid-age (25-55years) and 5 older adults (>55 years) (Hofmann et al., 2001). A third study of 12 subjects that reported no significant change in MM signals with age was not considered (Mader et al., 2002). We therefore set out to measure the MM spectrum in a structured cohort of healthy female and male adults aged between 20 and 70 years old, in order to characterize the age trajectory and sex dependence of the MM spectrum.

## 2. Methods

All data were acquired on a clinical 3T MRI scanner (Ingenia CX, Philips Healthcare, The Netherlands).

### 2.1 Preliminary investigations

In order to determine the optimal TI and TR for the metabolite-nulling experiments, a preliminary TI series acquisition was performed at two different TRs. PRESS-localized single-voxel spectra were acquired from two volunteers, acquiring 96 transients of 2048 datapoints per acquisition sampled at 2 kHz, 30 × 26 × 26 mm^3^ voxels localized in the centrum semiovale (CSO) and the posterior cingulate cortex (PCC) regions. For a TR of 3s, data were acquired with inversion times of 600 ms, 700 ms, 750 ms, 800 ms 850 ms, 900 ms, and 1000 ms. For a TR of 2s, data were acquired with inversion times of 400 ms, 500 ms, 550 ms, 600 ms 650 ms, 700 ms, and 800 ms. An adiabatic hyperbolic-secant inversion pulse with full-width half-maximum (FWHM) bandwidth 698 Hz was applied at 1.9 ppm for inversion. Philips-CHESS water suppression was re-optimized with an automated pre-scan routine for each TI.

### 2.2 Cohort

One hundred and two healthy volunteers were recruited with local IRB approval (Shandong University School of Medicine). The cohort was structured to include approximately ten male and ten female participants in each decade of age from the 20s, 30s, 40s, 50s, to the 60s. Exclusion criteria included contraindications for MRI and a history of neurological and psychiatric illness.

### 2.3 Acquisition

The acquisition protocol began with a T_1_-weighted MPRAGE scan (TR/TE/ 6.9/3.2 ms; FA 8°) with 1 mm^3^ isotropic resolution for voxel positioning and tissue segmentation. Metabolite-nulled MM spectra were then collected using PRESS localization (1.3 kHz refocusing bandwidth) with the above parameters except: TR/TE 2000/30 ms; 30 × 26 × 26 mm^3^ voxels localized in the CSO (predominantly white matter) and PCC (predominantly gray matter, Figure 1); and 600 ms inversion time (chosen based on preliminary studies – fuller discussions of results below); 96 transients sampled at 2 kHz. A slice-selective saturation pulse (20 mm thickness) was applied to suppress subcutaneous lipid adjacent to the voxel in CSO and PCC acquisitions. Water reference spectra were acquired without water suppression or pre-inversion.

**Figure 1:**
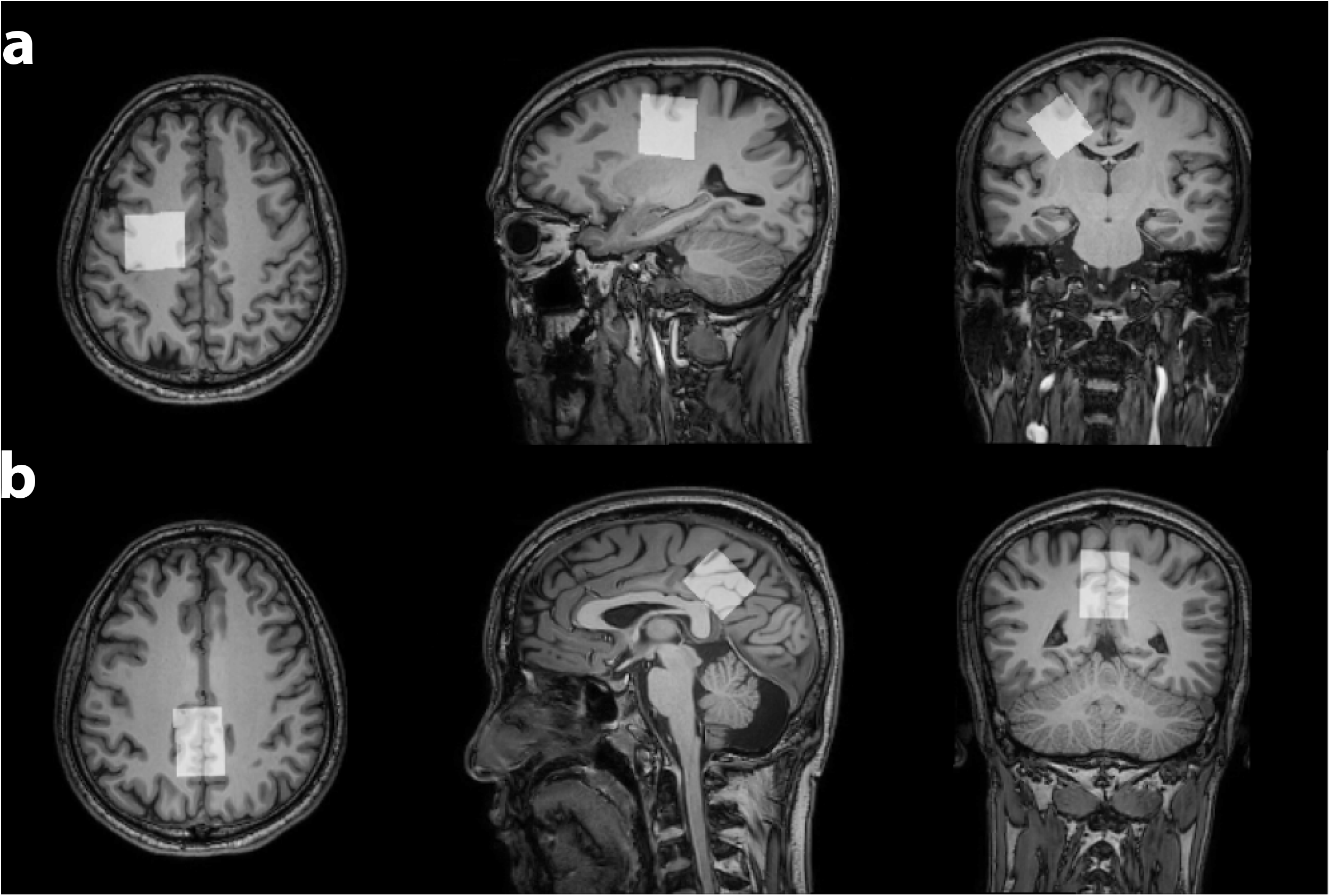
Voxel locations for (a) centrum semiovale (CSO) and (b) posterior cingulate cortex (PCC).

### 2.4 Analysis

#### 2.4.1 Metabolite residual cleaning and data selection

Spectra were processed using the Osprey software (Oeltzschner et al., 2020). Frequency-domain averaged metabolite-nulled spectra were modeled using a simulated reduced basis set (NAA, Cr, Cho, Glu) and a flexible spline baseline (0.1 ppm knot spacing) in order to model the residual metabolite signals. These modeled metabolite signals were then subtracted from the spectrum to yield a ‘clean’ MM spectrum (as illustrated in Figure 2). Individual-subject clean MM spectra were then sorted according to the standard deviation of the difference of the single-subject spectrum to the group-mean spectrum. An arbitrary threshold was chosen to exclude spectra that were substantially different from the mean, which resulted in exclusion of 3 and 6 spectra for the CSO and PCC regions, respectively. The major factor leading to exclusion was the appearance of out-of-voxel lipid signals.

**Figure 2:**
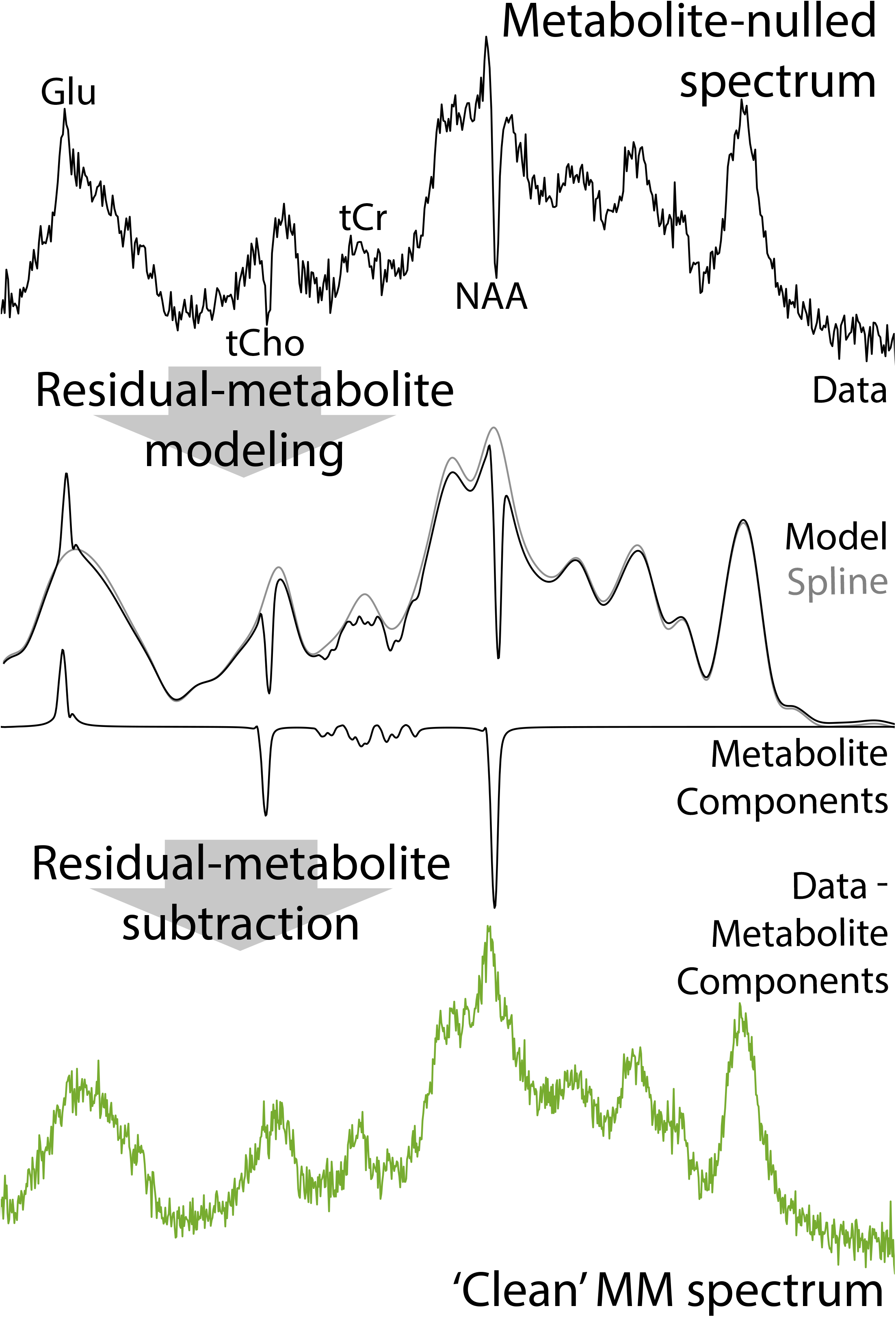
Osprey analysis workflow to remove residual metabolite signals and generate the ‘clean’ MM spectrum (shown in green). The spectrum is modeled with a flexible spline and residual metabolite signals; modeled metabolite signals are then subtracted from the data.

#### 2.4.2 Voxel segmentation and water referencing

Brain tissue segmentation was performed using Osprey with integrated voxel registration to the T_1_-weighted image and voxel segmentation (yielding relative tissue volume fractions f_GM_, f_WM_ and f_CSF_), based on functions implemented in SPM12 (Friston et al., 1994). Water reference spectra were modeled in the frequency domain as Gaussian-Lorentzian signals, and each subject’s MM spectrum was scaled by the water integral, adjusted for tissue-specific water visibility and relaxation based on literature values (Wansapura et al., 1999).

#### 2.4.3 MM peaks modelling

MM spectra were then averaged and modeled as a sum of 14 Gaussian peaks between 0.5 and 4.1 ppm. The MM peaks included were M_0.94_, M_1.22_, M_1.43_, M_1.69_, M_2.04_, M_2.27_, M_2.66_, M_3.01_, M_3.21_, M_3.71_, M_3.79_, M_3.87_, M_3.97_ and M_4.20_, where the respective subscripts indicate the chemical shift in ppm of the MM peaks. Model parameters included the width and amplitude of each Gaussian (2 × 14 = 28 parameters), and three additional parameters modeled a global frequency shift (to account for small differences of ppm referencing) and a baseline offset and linear baseline slope (for a total of 31 parameters). Optimal model parameters were determined using non-linear leastsquares optimization in MATLAB (R2020b, MathWorks, Natick, USA). The cohort-average spectrum was modeled first; these parameters were then used to initialize modeling of the individual-subject spectra. For each subject, modeled MM signal areas were extracted for each Gaussian peak; for display purposes, the integrals of M_3.71_, M_3.79_, M_3.87_ and M_3.97_ were combined to accommodate imperfect resolution in the spectrum at 3T.

### 2.5 Statistical Analysis

For each of the 14 MM peaks, Pearson’s correlation coefficients were calculated to investigate relationships between age and MM signal integral. Between-age-group (5 groups: 20s, 30s, 40s, 50s, 60s) differences were assessed using one-way ANOVA. Differences between sexes were assessed using independent-samples t-test. Correlations and differences between segmented brain tissues (CSF, GM and WM) due to age and sex groups were analyzed using Pearson’s coefficient and independent t-test respectively. All statistical analyses were performed using R (Version 4.0.2) in RStudio (Version 1.2.5019, Integrated Development for R. RStudio, PBC, Boston, MA). Holm-Bonferroni correction (with an adjusted p-value threshold of 0.004) was used to account for statistical testing of the 14 MM peaks. This correction does not apply to correlations between brain tissue fraction and age.

## 3. Results

TI series acquired at TR of 2s and 3s are shown in Figure 3a and 3b respectively. As expected, the metabolite nulling time is longer for the 3s data than the 2s data, due to more complete relaxation toward equilibrium before inversion. Null times for Cho, Cr_3.0_ and NAA singlets respectively are ~[600 ms, 650 ms, 700 ms] at TR 2s and ~[750 ms, 800 ms, 850 ms] at TR 3s. Note that these values are dependent on metabolite T_1_ (and therefore brain region) and inversion efficiency. Given that the coupled metabolite T_1_s are likely to be closest to Cho, and singlet signals are easier to remove in post-processing, we resolved to use the Cho null-point (600 ms) for the cohort acquisition. At the respective Cho-nulling TI, residual NAA and Cr signals in the TR 2s and TR 3s spectra look very similar, so TR 2s was chosen to deliver better temporal SNR.

**Figure 3:**
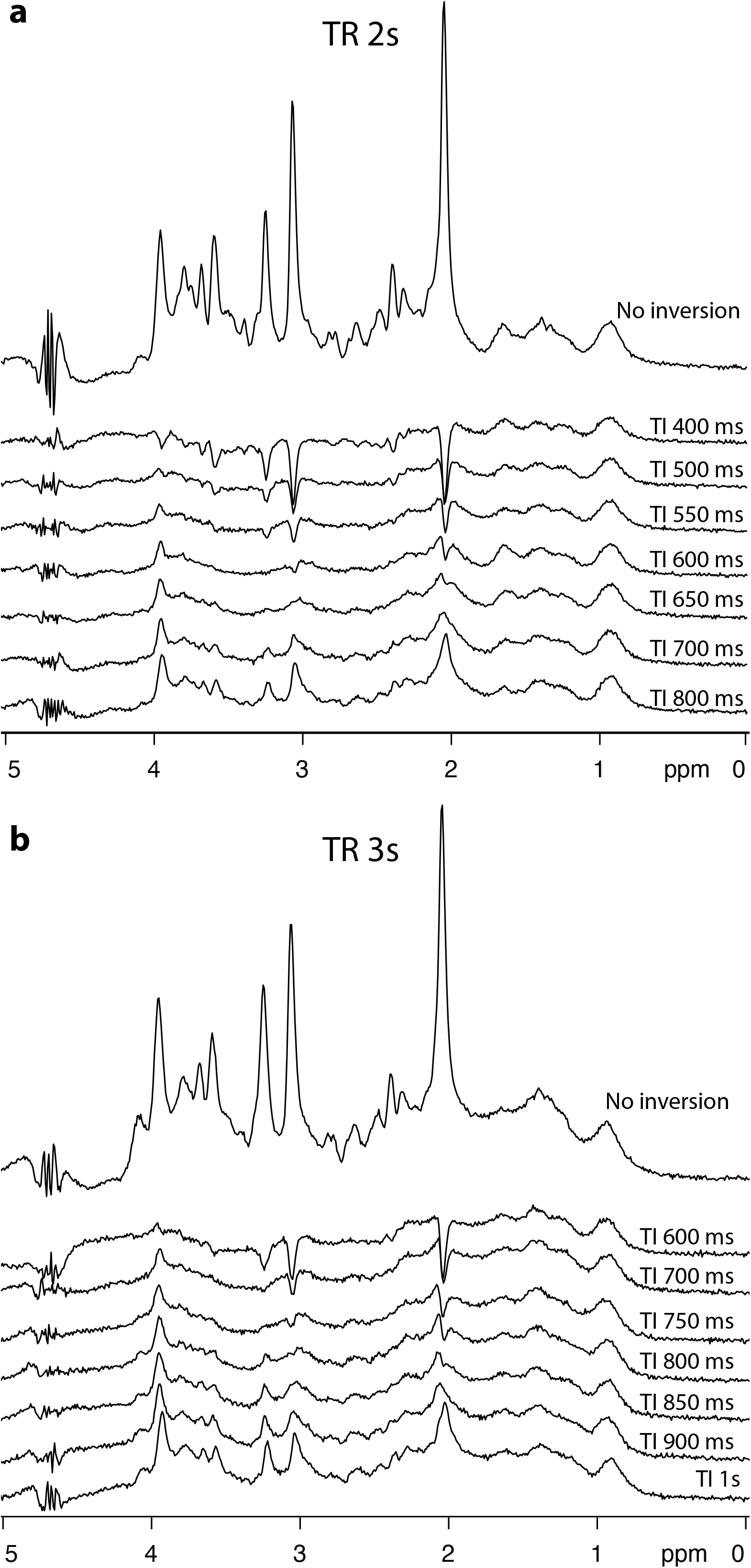
Preliminary experiments establishing the Cho-nulling TI at TR 2s (a) and TR 3s (b). MM signals are relatively consistent, while metabolite signals are inverted at short TIs, pass through a null point and are positive at long TIs. The choline null-points are: (a) 600 ms and (b) 750 ms.

99 spectra for CSO and 96 for PCC were included in further analysis as shown in Figure 4a, with the demographic groupings outlined in Table 1. Average water linewidths in CSO and PCC were both 6.6 Hz, indicating excellent data quality. The MM signals used to model the spectra with labels M_0.94_-M_4.20_ are shown in Figure 4b. Average spectra are shown for male and female participants in Figure 4c, illustrating good concordance. Spectra averaged from each decade of life are shown in Figure 4d. Note that the agreement between spectra is greater between 2 and 4 ppm than between 1 and 2 ppm. Pearson’s correlation coefficients suggested no significant relationships between age and MM integrals (all p > 0.06) as illustrated in Figure 5 and one-way ANOVA indicated no significant between-age-group differences (all p > 0.07). No significant differences were found between male and female participants (all p > 0.03).

**Figure 4:**
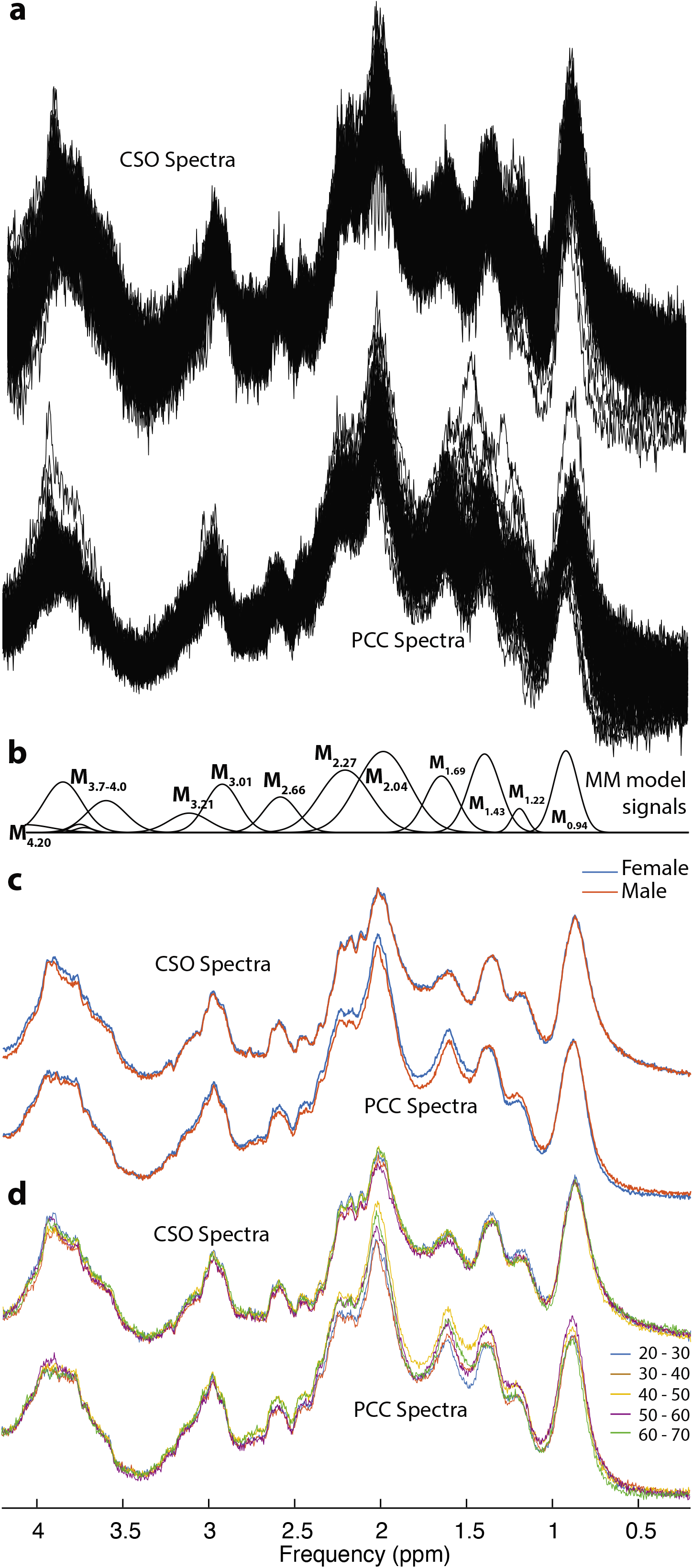
a) MM spectra from 99 and 96 subjects are overlaid for CSO and PCC respectively. b) The MM model signals. c) The mean spectra for male and female subjects. d) The mean spectra for each decade of age.

**Figure 5:**
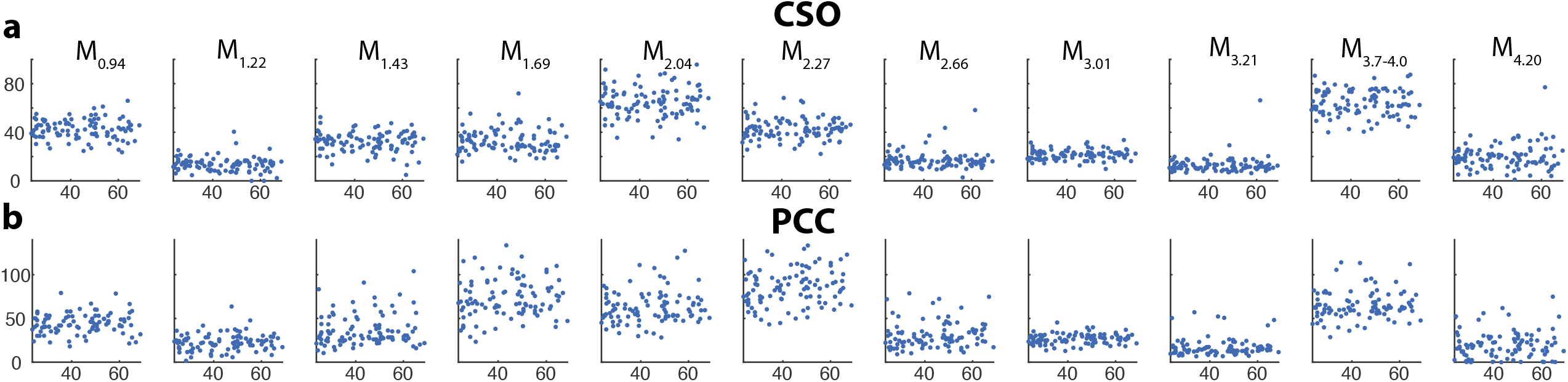
Correlations between water-scaled MM signal integrals and age for (a) CSO and (b) PCC. MM signals are labeled by chemical shift as per Figure 4b. The x-axis shows age in years and the y-axis water-scaled MM signal amplitudes. None of the plots show a statistically significant correlation (p > 0.06 for all tests).

**Table 1:**
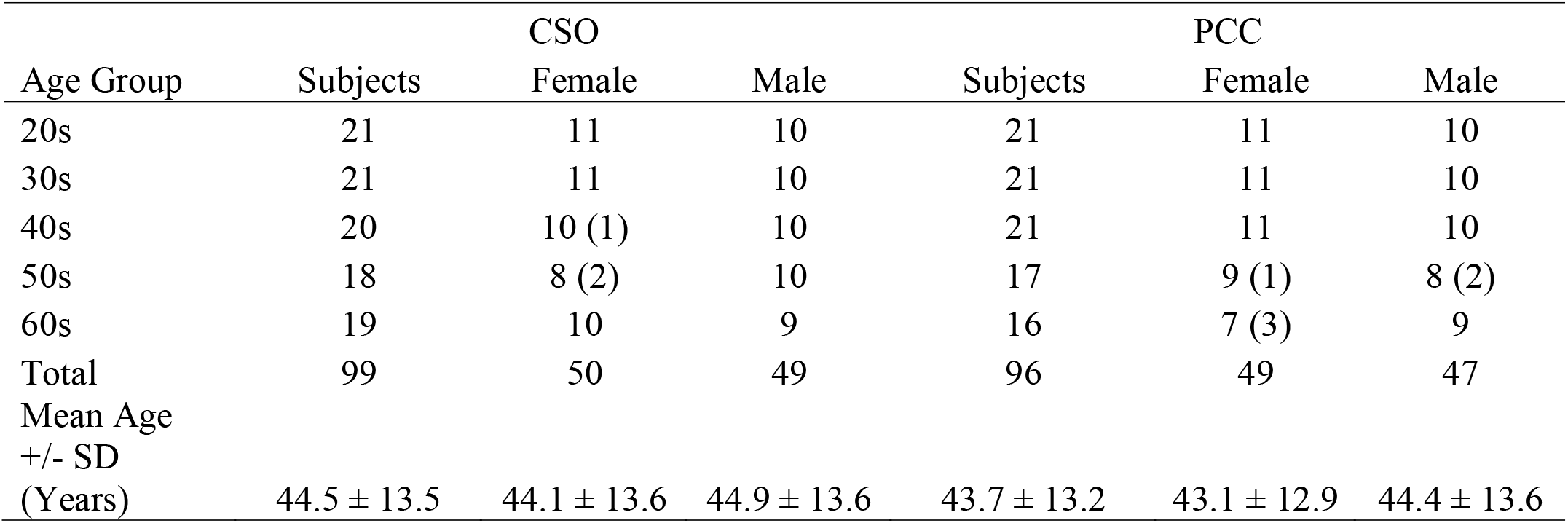
Description of the cohort. Numbers of subjects in each demographic group are shown after excluding low-quality datasets (numbers of spectra excluded in each group are shown in parentheses).

The voxel tissue content was segmented as 79±4% WM, 19±4% GM, and 2±1% CSF for CSO and 28±3% WM, 61±4% GM and 11±4% CSF for PCC. In CSO, correlations between tissue fraction and age indicated a slow but significant reduction in the predominant WM voxel fraction as a function of age, accounted for in equal amounts by a significant increase in CSF (Figure 6a). In PCC, a significant decrease was observed in the predominant GM voxel fraction as a function of age while both WM and CSF increased with age (Figure 6b). CSF fraction was significantly higher in male subjects than in female subjects (p < 0.05 for both CSO and PCC).

**Figure 6:**
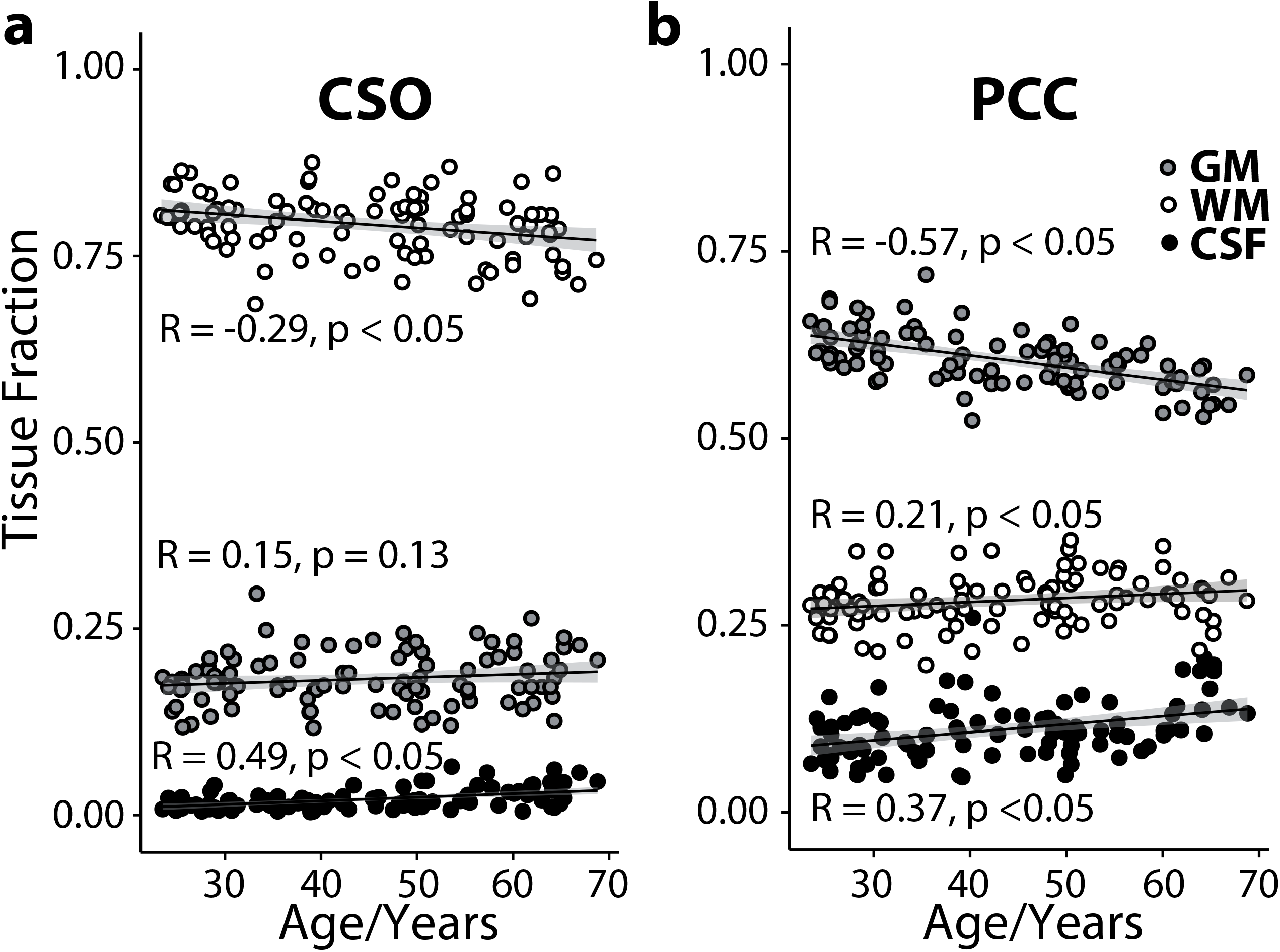
Voxel segmentation as a function of age for (a) CSO and (b) PCC.

## 4. Discussion

Macromolecular spectra were successfully acquired in the CSO for 99 subjects and in the PCC for 96 subjects, using pre-inversion to null the metabolite signals. Due to the range of metabolite T_1_s, additional post-processing was performed to remove residual metabolite signals from the spectra, a consensus process (Cudalbu et al., 2021) implemented within Osprey (Oeltzschner et al., 2020). These ‘clean’ MM spectra did not differ between male and female subjects, or demonstrate a relationship with age. The MM data acquired in this study will be integrated as an option in Osprey to be used for linear-combination modeling of short-TE spectra and be available on NITRC for further analysis. The MM data acquired in this study will be integrated as an option in Osprey to be used for linear-combination modeling of short-TE spectra.

The results of this study differ from previous reports of age-related changes in specific peaks in the MM spectrum (Hofmann et al., 2001; Marjanska et al., 2018). These two studies were not mutually consistent – the larger study failing to show significant differences between an older group and either the younger, or mid-age groups. A third prior manuscript suggests there may be age-related change only in the M_0.9_ signal, but recognizes that this does not meet significance after multiple comparison correction (Mader et al., 2002). It is possible that the differences between these studies are explained by the different brain regions studied or other acquisition differences. The water referencing procedure for relaxation correction also differs between the three studies. The CHESS water suppression applied in our study with 140 Hz bandwidth has a substantial suppression effect on the M_4.20_ peak. In Marjanska et al. (Marjanska et al., 2018), VAPOR was implemented but the bandwidth of the pulse was not specified. A reduction of the bandwidth to below 80 Hz would largely avoid any unintended signal suppression on M_4.20_. We are unaware of any other previous reports of a relationship between M_4.20_ and age or sex. Our data emphasize the importance of considering age as a continuous, rather than categorical, variable. There is an on-going need for tissue water and metabolite relaxation reference datasets throughout the lifespan. Without that, any quantitative study of aging is hard to interpret.

It is interesting to note that the spectra excluded from this study, 3 for CSO and 6 for PCC, had unacceptably high levels of lipid signals in the spectrum, associated with imperfect localization of signal to the prescribed volume. These signals appear in the same region of the spectrum that is most variable in Figure 4d and in which significant age differences have been shown in previously mentioned studies of MM in the context of aging (Hofmann et al., 2001; Marjanska et al., 2018). The largest MM signal variability can be observed in PCC from M_0.94_ to M_2.27_ as shown in Figure 4d. This range is known to have 3 major lipid signals, at 0.9, 1.3 and 2.1 ppm. Unintentional excitation of out-of-voxel lipid signal (which is greater for the PCC location) likely explains the larger variance. Although data with the greatest lipid signal were excluded, effects from residual signals remain visible. An exploratory analysis applying a substantially stricter exclusion criteria to our dataset does not alter the major findings (not reported here).

Significant relationships were observed between age and brain tissue segmentation, consistent with previous studies (Ge et al., 2002; Giorgio et al., 2010). It is possible that these represent a combination of real changes in tissue volume and the interaction between changes in T_1_-weighted image contrast and the segmentation routine. In both regions of our results, GM-rich PCC and WM-rich CSO, the major tissue fraction reduces with age, consistent with brain atrophy due to normal aging, as widely studied (Guttmann et al., 1998; Kitagaki et al., 2000; Meier-Ruge et al., 1992). The observation, that in both regions the minor tissue fraction (i.e. GM in CSO, or WM in PCC) tends to increase with age, is consistent with a loss of image contrast, and segmentation reverting toward the mean. Prior studies suggested that white matter atrophy could be an indirect indicator of nerve cell loss, volume changes have been characterized as quadratic or linear (Farokhian et al., 2017; Meier-Ruge et al., 1992). In our study, male subjects had significantly higher CSF compared with female subjects in both CSO and PCC regions, possibly reflecting accelerated aging (May et al., 1990).

The acquisition of metabolite-nulled spectra to characterize the MM spectrum is a technically challenging undertaking. Ideally, such an experiment should simultaneously null all metabolite signals, while leaving MM signals fully relaxed. Both single- and double-inversion approaches have been applied to achieve this, with double inversion giving better metabolite nulling across a range of TIs at the expense of greater T_1_-weighting of MM signals. In this study, a single inversion pulse of bandwidth 698 Hz (99% inversion) was applied at an offset of −356 Hz from the water resonance (i.e at 1.9 ppm), so as to invert the entire spectral range of interest from - 0.85 to 4.65 ppm. This permits high inversion efficiency for the metabolite signals, while minimizing perturbation of the water signal. The selection of TI was based on preliminary testing in several subjects, in which the different T_1_ times of the metabolites indicated a different null point for each. In this study, the Cho null-point at 600 ms was chosen which inevitably led to small, but noticeable, amounts of residual NAA and Cr signal. Although a consensus-recommended ‘cleaning’ procedure was used to model and subtract out these residual signals, the procedure was not 100% successful in all subjects. Even in the absence of residual metabolite signals, modeling the relatively featureless MM spectra with a large number of overlapping Gaussian signals is itself challenging, and there is likely substantial covariance between signal amplitude and baseline terms. Future studies will be able to refine the analysis procedure using the publicly available data from this study.

It has previously been reported that MM T_1_ relaxation times were significantly different between GM- and WM-rich voxels. M_3.75_ (labeled as M_3.71_ and M_3.79_ in our study) and M_3.21_ had the greatest difference in T_1_-weighting between GM and WM (Murali-Manohar et al., 2021). The same study group also reported T_2_ relaxation times of MMs, with smaller differences between GM and WM (Murali-Manohar et al., 2020). The effective T_2_ varies substantially between MM signals at 3T (Landheer et al., 2020). Tissue corrections for MM relaxation times have not been applied in this manuscript because T_1_ and T_2_ relaxation times have not yet been reported as a function of age.

In conclusion, this manuscript presents MM spectra acquired in a large, structured crosssectional cohort to investigate MM changes with age and sex. In contrast to some of the prior literature, no significant age-related changes are observed. There is much value in resolving the question of whether the MM spectrum changes with age and other factors including different tissue regions (WM and GM), as it has strong implications for appropriate analysis workflow. These results support the use of averaged MM spectra in linear-combination modeling for many applications. However, given the diverse MM relaxation behavior, it is important to match the TE of MM reference spectra and data.

## Acknowledgements

This work was supported by NIH grants R01 EB016089, R01 EB023963, K99/R00 AG062230, K99 DA051315, P41 EB015909, P41 EB031771, KO1 AA025306, and S10 OD021648. This study was also supported by Natural Science Foundation of Shandong (Grant No. ZR2020QH267).

